# The astrocyte-enriched gene *CG11000* plays a crucial role in the development, locomotion and lifespan of *D. melanogaster*

**DOI:** 10.1101/839811

**Authors:** Hadi Najafi, Kyle Wong, Ammar Salkini, Hongyu Miao, Woo Jae Kim

## Abstract

The brain of *Drosophila melanogaster* is a complex organ with various cell types, orchestrating the physiology and behaviors of the fly. While each cell type in the *Drosophila* brain is known to express a unique set of genes, their complete genetic profile is still unknown. Advances in the RNA-sequencing techniques at single cell resolution facilitate identifying novel cell type-specific markers and/or examining the specificity of the available markers.

In this study, exploiting a single cell RNA sequencing data of *Drosophila* optic lobe (which comprises two thirds of the brain with extensive cell type diversity), we first categorized cell types based on their known molecular markers, then the genes with enriched expression in astrocytes were identified. Consistent with previous findings, the known glial markers *CG34335*, *Inx2* and *nrv2* as well as the astrocytic genes *CG9394*, *Eaat1*, *Gat*, *Gs2* and *CG1552* exhibited enriched expression in the identified astrocyte cluster. Moreover, we identified *CG11000* as a gene with positive expression correlation with the astrocytic marker *Eaat1*. The positive expression correlation between *CG11000* and *Eaat1* genes was also observed in the single-cell RNA-sequencing data of *Drosophila* mid-brain as well as in the bulk RNA-sequencing data of *Drosophila* whole brain during development.

Immunostaining of the brains dissected from adult flies showed overlapping fluorescence signals of *CG11000* and *Eaat1* expression, supporting co-expression of these genes in a set of single cells in *Drosophila* optic lobe. At the physiological level, RNAi-mediated suppression of *CG11000* impeded the normal development of male flies without any effects on females. In adult flies, *CG11000* suppression affected the locomotion activity and lifespan of *D. melanogaster* in an astrocyte-specific manner, suggesting pivotal role of *CG11000* gene in astrocytes.

## Introduction

The central nervous system of *Drosophila melanogaster* is a powerful model system to study development [1–3], physiology [4–6] and behaviors [7, 8] of other advanced organisms (including human), due to the great homologies between the cellular and molecular composition of the *Drosophila* nervous system and that of other organisms [9–12]. The *Drosophila* brain is composed of various cell types which are primarily characterized by their morphology, spatial organization and limited number of genetic driver lines (e.g., Gal4/UAS) [13–15], hence necessitate studying this organ deeper at single cell resolution.

In neuroscience, several classical cell type markers are commonly used to indicate and/or target a particular population of cells, considered to be the cells of a single type [16, 17]. At least seven main types of neural cells were identified in *D. melanogaster*, each characterized by the expression of a unique set of genes known as their molecular markers [16]. These cell types include perineurial glia (PNG), sub-perineurial glia (SPG), cortex glia (CG), astrocyte-like glia (ALG or astrocytes), neuropile ensheathing glia (EGN), tract ensheathing glia (EGT) and neurons (Neu), each possesses its own genetic driver lines [16].

Since the expression pattern of a single or a set of multiple genes could not represent the full genetic profile of a cell type [18, 19], additional analyses are required to characterize them. Also, besides the previously-known molecular markers for the respective cell types, some novel genes may exist in *D. melanogaster* such that a specific cell type can be discerned.

With the advent of single cell RNA sequencing (scRNA-seq) technologies, analysis of the entire transcriptome of an individual cell is feasible which facilitates determination of cellular identities in all multicellular organisms including *D. melanogaster*. By providing the full genetic profile of an individual cell, scRNA-seq is a convenient strategy for identification of novel cell types [20–22] and also reexamining the specificity of the pre-existing molecular markers for each cell type of *D. melanogaster* [23, 24].

In this study, we will take the advantages of the availability of scRNA-seq data of *Drosophila* brain to perform clustering of the sequenced single cells based on their expression pattern for the previously-known molecular markers, then finding novel genes with differential expression pattern across the clusters. Identification and functional characterization of the genes with enriched expression in a particular cell type in *D. melanogaster* could provide a new insight into the mechanistic role of the respective cell type.

## Materials and methods

### Datasets

A single cell RNA-sequencing (scRNA-seq) data belonging to the *Drosophila* optic lobe was obtained from the NCBI Gene Expression Omnibus (GEO) database (accession number: GSE103771), which covered the expression profile of 17,272 *Drosophila* genes in a population of 120,000 single cells from optic lobe [25].

Another single cell RNA-sequencing (scRNA-seq) data was from the *Drosophila* mid-brain, and was obtained from the NCBI Sequence Read Archive (SRA) database (accession number: SRP128516), which covered the expression profile of 12,868 *Drosophila* genes in a population of 28,695 single cells from mid-brain [26].

Two bulk RNA-sequencing data of *Drosophila* whole brain were obtained from the NCBI GEO: one contained the expression profile of all *D. melanogaster* genes across four different developmental stages (days 5, 20, 30 and 40 post-eclosion; accession number: GSE107049) [27] and another one contained the expression profile of all *D. melanogaster* genes across three different developmental stages of the fly (days 3, 7 and 14 post-eclosion; accession number: GSE199164).

### Bioinformatics analyses

In order to have a set of clusters comprising the main neural cell types of *D. melanogaster* (i.e., perineural glia (PNG), subperineural glia (SPG), cortex glia (CG), neuropil ensheathing glia (EGN), tract ensheathing glia (EGT), astrocyte-like glia (ALG) and neurons), first, the expression matrix belonging to the scRNA-seq data of *Drosophila* optic lobe in the study of Konstantinides et al. (accession number: GSE103771) [25] was downloaded and used for downstream analyses. Then, this matrix was analyzed in accordance with the expression pattern of the previously-known *Drosophila* marker genes specific to PNG, SPG, CG, EGN, EGT, ALG and neuronal cell types and the single cells expressing the markers of a respective cell type and negative for the expression of the markers of other cell types were identified and presented as a heatmap of expression level using the online tool Heatmapper [28] with color codes (red and black colors refer to positive and negative expression for the markers, respectively).

To confirm the above-mentioned cell type classification, hierarchical clustering was performed in accordance with the similarities between the identified single cells based on their transcriptome profile and expressed as a heatmap with color codes for the Pearson correlation coefficient, using the ggplot2 [29] and pheatmap [30] packages in R. Also, principal component analysis (PCA) was performed to distinguish the cells/clusters with distinct transcriptome profile, using R program.

The lists of differentially expressed genes (DEGs) between clusters of interest were identified by the R package limma [31], considering the log_2_fold-change (log_2_FC) ≥ 1 and *P*-value < 0.05 as the significance cutoff.

To confirm the differential expression pattern of the candidate genes in a particular cell type/cluster, the unpaired t-test method [32] was applied considering adjusted *P*-values (Benjamini–Hochberg correction) < 0.01 and log_2_FC ≥ 1 as the significance cutoff. Venn diagram was performed for obtaining the list of common DEGs between the examined clusters, using the web-based tool Venny [33]. Pairwise alignment between the transcripts of *CR34335* and *CR40469* genes was performed in ClustalW [34, 35].

### Fly stocks and crosses

All the *D. melanogaster* lines in this study were obtained from the Bloomington Drosophila Stock Center (BDSC). After being subjected to stocks, they were cultured in standard conditions (standard cornmeal-agar yeast fly food at 25 °C in a 12h light:12h dark condition) following the previous description [36]. CO_2_ served as an anesthetic.

For the crosses of fluorescence labeling experiments, the two binary systems Gal4/UAS- mCD8RFP and LexA/LexAop-mCD8GFP were employed simultaneously in individual male and female flies. The ALG cells of *Drosophila* CNS were labelled in accordance with the expression of a membrane-bound GFP (mCD8GFP) under the control of *Eaat1* gene promoter, fused to the sequence of LexA (*Eaat1*-LexA) (BDSC#: 52719).

A transgenic fly line carrying the *CG11000*-Gal4 was used to drive the expression of a membrane-bound RFP (mCD8RFP) in progenies (both in female and male) under the control of *CG11000* gene promoter (BDSC#: 72713). Therefore, the expression pattern of RFP will reflect the pattern of *CG11000* promoter activity across the *Drosophila* single cells. Table 1 lists the fly stocks and the crossing scheme for immunostaining experiment.

**Table 1.** Crossing scheme for imaging experiments of the *Drosophila* brain. The line A contains RFP and GFP reporter genes under the control of UAS and LexAop sequences, respectively. Line B expresses LexA protein in an ALG-specific manner driven by *Eaat1* promoter. The fly line C (of F_1_ generation) was generated by crossing the lines A and B (parents), and then it was crossed to the *CG11000*-Gal4 line (i.e., D) to produce F_2_ generation. Among the flies of F_2_ generation, the males and females that lacked balancer chromosomes but contain the complete components of the two binary systems UAS/Gal4 and LexA/LexAop were selected for brain dissection. The females (♀) of all crosses were virgin.

| Lines | Fly stocks (genotype & BDSC ID) | Cross scheme |
| --- | --- | --- |
| A | UAS-mCD8RFP, lexAop-mCD8GFP ; Sp/CyO ; TM2/TM6B (# 32229) | **P:** A ♀ × B ♂ |
| B | w<sup>-</sup> ; *Eaat1*-LexA/CyO ; TM2/TM6B (#52719) |  |
| C | UAS-mCD8RFP, lexAop-mCD8GFP ; ALG-LexA/CyO ; TM2/TM6B<br>(This study) |  |
| D | w<sup>-</sup> ; +/+ ; *CG11000*-Gal4 (#72713) |  |
| E | UAS-mCD8RFP, lexAop-mCD8GFP ; ALG-LexA/+ ; TM6B/<br>*CG11000*-Gal4 (This study) |  |

For functional analysis of the genes, the Gal4/UAS-RNAi system [37] was employed to deplete the function of genes of interest in a cell type-specific manner. Table 2 summarizes the fly stocks (parents) together with their crossing schemes, ensuring progenies to express the *CG11000* RNAi in a cell type-specific manner. In detail, the virgin females transgenic for the UAS-*CG11000* RNAi construct (BDSC#: 51918) were crossed to different male driver lines expressing Gal4 under the control of the promoter of a set of cell type-specific genes (table 2). The promoter of *shn* gene drives Gal4 expression in PNG, *Mdr65* in SPG, *wrapper* in CG, *CG9657* in EGN, *CG34340* (*Drgx*) in EGT, *CG5264* in ALG, *repo* in all glia, and *nSyb* drives Gal4 expression in neurons [16, 23, 38]. After crossing the flies, the progenies with complete components of the Gal4/UAS binary system (i.e., x-Gal4 > UAS-CG11000 RNAi) were collected based on their phenotypes (table 2) and functional analyses of *CG11000* gene were performed for them. As the control group of the experiment, flies without expression of *CG11000* RNAi but with the same genetic background were used. In fact, the flies of the control group contained the UAS- *CG11000* RNAi construct in their genome but lacked any expression of Gal4 proteins. Thus, no functional RNAi will be produced in this group.

**Table 2.** Summary of the crosses and experimental genotypes for behavioral analyses. To generate the transgenic flies with expression of *CG11000*-RNAi in cell type-specific manner, the male flies of different Gal4 driver lines were crossed to the females harboring UAS-*CG11000*-RNAi in their genome (BDSC#: 51918). All the females (♀) for the above-mentioned crosses were virgin. Among the progenies of the above-described crosses, the flies with proper genotypes (harboring both cell type-specific expression of Gal4 and UAS-*CG11000* RNAi) were selected based on the absence of balancer chromosomes (e.g., CyO, TM3 and TM6B).

| Parents of the crosses (genotype & BDSC ID) |  | Progeny phenotypes<br>(♀ and ♂) | RNAi-expressing cell type |
| --- | --- | --- | --- |
| UAS-RNAi transgenic line<br>(♀) | Gal4 driver lines (♂) |  |  |
| w<sup>-</sup>; +/+; UAS-*CG11000*-RNAi<br>(# 51918) | w<sup>1118</sup>; +/+; +/+ | wild type | - |
|  | w<sup>1118</sup>; +/+; shn-GAL4/TM3 (#40436) | Non-TM3 | PNG |
|  | w<sup>1118</sup>; +/+; Mdr65-GAL4/TM3 (#50472) | Non-TM3 | SPG |
|  | w<sup>1118</sup>; +/+; CG-GAL4/TM3 (#45784) | Non-TM3 | CG |
|  | w<sup>1118</sup>; +/+; CG9657-GAL4/TM3 (#39157) | Non-TM3 | EGN |
|  | w<sup>1118</sup>; +/+; Drgx-GAL4/TM3 (#39908) | Non-TM3 | EGT |
|  | w<sup>1118</sup>; +/+; btn-GAL4/TM6B (#45914) | Non-TM6B | ALG |
|  | w<sup>1118</sup>; +/+; nSyb-GAL4/TM6B (#68222) | Non-TM6B | Neurons |

### Immunostaining and imaging

Immunostaining was performed according to the previously-reported protocols [39, 40]. In brief, brains were dissected from adults 5 days after eclosion, fixed with 4% formaldehyde for 30 min at room temperature, washed with 1X PBST (1X PBS containing 0.5% Triton-X100) three times (30 min each) and blocked in 5% normal donkey serum for 30 min. The brains were then incubated with primary antibodies in 1X PBST at 4 °C overnight, followed by incubation with fluorophore-conjugated secondary antibodies for 1 hour at room temperature. Subsequently, brains were mounted with antifade mounting solution (Invitrogen, catalog number S2828) on slides for imaging.

The primary antibodies were chicken polyclonal anti-GFP (Aves Labs, catalogue number GFP- 1010, 1:1000 diluted), rabbit polyclonal anti-RFP (Invitrogen, catalogue number 710530, 1:250 diluted) and mouse monoclonal anti-Bruchpilot (nc82) (DSHB, ID AB-2314866, 1:50 diluted). The applied fluorophore-conjugated secondary antibodies comprised Alexa Fluor 488-conjugated goat anti-chicken (Invitrogen, catalogue number A32931, 1:100 diluted), RRX-conjugated donkey anti-rabbit (Jackson Laboratory, catalogue number AB-2340613, 1:100 diluted), and Dylight 405-conjugated donkey anti-mouse (Jackson Laboratory, catalogue number AB- 2340839, 1:100 diluted). Fluorescence imaging with maximum projections of z-stacks were performed under a confocal laser scanning microscope (Zeiss LSM 7 MP) and the acquired images were processed using the NIH ImageJ [41], and presented by Adobe Illustrator CC 2023 for Mac.

### Developmental assay

To assess the possible function of *CG11000* gene in the development of *D. melanogaster* from pupae to adulthood stage, the newly eclosed *D. melanogaster* of the crosses in table 2, were collected and counted based on their genotypes. After counting, data were presented as the percentage of male and female flies expressing *CG11000* RNAi to the total number of eclosed flies.

### Climbing assay

For climbing assay, flies were separated into groups of 25 adults per each vial and kept at 29 °C. At the time of experiment, the groups of 25 flies were placed in an empty climbing vial and then tapped down to the bottom. They were allowed to climb from the bottom to top of the vial (8 cm) for 20 sec. To evaluate the climbing pattern of each genotype, the climbing height achieved by each genotype was calculated in every 5 sec after the tapping and initiating negative geotaxis. The results of climbing assay were obtained through video recording of climbing flies and then processed using ImageJ software [42]. The data are expressed as violon plots using Prism version 10 (GraphPad Software, San Diego, CA, USA). Flies lacking any expression of CG11000 RNAi but with the same genetic background were used as control.

### Lifespan assay

Lifespan was measured at room temperature according to the standard protocols [43]. In brief, newly ecloded flies (0 to 3 days) were collected (50 per genotype) and then placed in two vials (25 flies per vial), and transferred to fresh vials every two days. Survival was recorded for each genotype (sum of two vials). We scored flies stacked in the food as death events in all the vials analyzed. The survival curves was created with Prism version 10 (GraphPad Software, San Diego, CA, USA) using the method of Kaplan and Meier [44]. For statistical significance of the experiment, one-way ANOVA method was used and the death rate of each genotype was compared to control group which contained the flies without expression of CG11000 RNAi but with the same genetic background. *P*-values < 0.01 indicated the differences that achieved statistical significance.

### Statistical analysis

Statistical analysis and graphic representations were conducted using Prism 10 for Mac (GraphPad Software, San Diego, CA, USA). Unpaired t-test or one-way ANOVA (using the Bonferroni’s multiple comparison) were applied, which was dependent on the measurements analyzed in the corresponding experiment. The averages with ± SEM in all cases were plotted. *P*- values were determined through two-tailed unpaired Student’s t-test, unless otherwise specified, using GraphPad Prism 10 software. ∗*P* < 0.05; ∗∗*P* < 0.01; ∗∗∗*P* < 0.001 and ∗∗∗∗*P* < 0.0001.

## Results

### Cell type identification of the single cells in *Drosophila* optic lobe based on known molecular markers

Each cell type of the *Drosophila* central nervous system is characterized by expression of a unique set of genes, known as their molecular markers [15, 16, 45]. We used these markers (Fig. 1A) to identify the type of the single cells in *Drosophila* optic lobe, sequenced by Konstantinides et al. (GEO accession number: GSE103771; n = 120,000 cells) [25], so that a particular cell type is positive for expression of its own markers while negative for the markers of other cell types. According to these parameters, 156 single cells were identified as neurons, being positive for *elav* and *nSyb* expression while negative for all glial markers, 80 single cells were identified as astrocyte-like glia (ALG or astrocytes), expressing five known astrocytic markers (e.g., *Eaat1*, alrm, *Gs2*, *ebony* and *Gat*) and lacking the expression of other glial and neuronal markers. Also, expression analysis of the markers specific to PNG (*CG4797*, *shn* and *gem*), SPG (*Mdr65* and *moody*), CG (*Cyp4g15* and *wrapper*) and EG (*CG9657* and *CG34340*) has identified 56 single cells belonging to these glial cell types other than astrocytes (assigned as non-ALG glia) (Fig. 1B).

**Figure 1.**
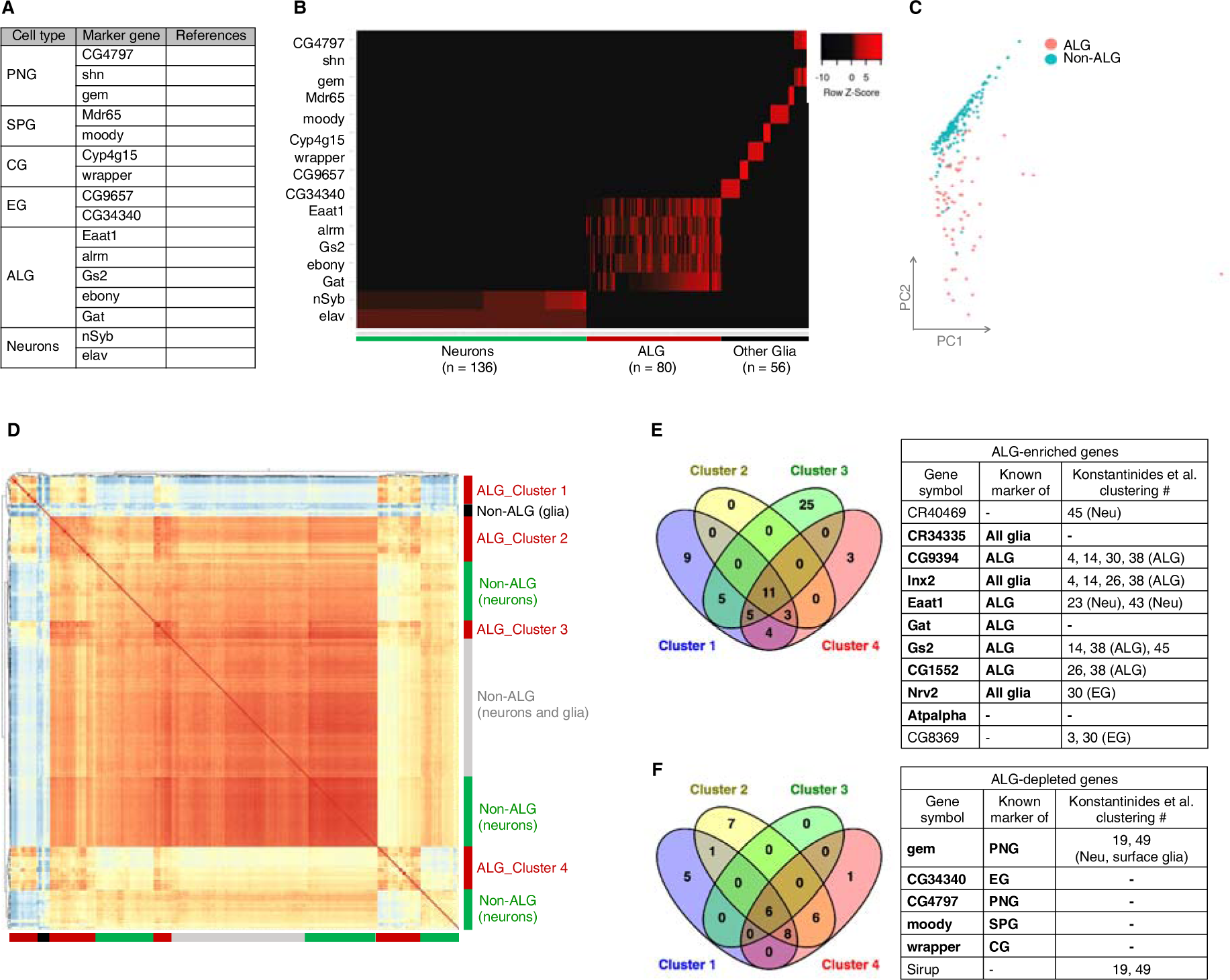
Cell type identification of the sequenced single cells in *Drosophila* optic lobe. **A)** The list of well-characterized markers in *D. melanogaster* nervous system, used in this study for cell type identification. Perineurial glia (PNG) are characterized by expression of *CG4797*, *shn* and *gem* genes; sub- perineurial glia (SPG) by *Mdr65* and *moody* genes; CG by *Cyp4g15* and *wrapper* genes; EG by *CG9657* and *CG34340* genes; ALG by *Eaat1*, *alrm*, *Gs2*, *ebony* and *Gat* genes; and neurons are characterized by the expression of *nSyb* and *elav* genes. (References in the table will be added). **B)** The classified single cells of *D. melanogaster* optic lobe based on the expression pattern of the selected markers. Red color denotes ‘expression’ while black color denotes ‘lack of expression’ for each marker. A row Z-score was used to represent the expression level of the markers. **C)** The principal component analysis (PCA) plot distinguishes the astrocyte-like glia (labeled as ALG; red dots) from other glial and neuronal cells (labeled as non-ALG; green dots) based on the entire transcriptome profile of the cells. **D)** Hierarchical clustering method categorized ALG cells into four different clusters (clusters 1 - 4), all exhibiting distinct global transcriptome profile form other cell types (i.e., non-ALG glia and neurons). **E)** Venn diagram illustrates the number of the genes upregulated in each ALG cluster. The table lists the common upregulated genes shared between all ALG clusters. Bold-face fonts represent the genes with cell type-specificity reports in previous studies. If the genes were previously attributed to a particular cell type by Konstantinides et al. [25], the features of their clusters are presented in the table for comparison. F) Venn diagram illustrates the number of the genes downregulated in each ALG cluster. The table lists the common downregulated genes shared between all ALG clusters. Also, the clustering information of Konstantinides et al. study is presented for each gene.

Thereafter, global transcriptome profile of the identified cells was examined through principal component analysis (PCA) and results showed a distinctive transcriptome profile for ALG, compared to the non-ALG glia and neurons (non-ALG cluster) (Fig. 1C). Further classification of the identified single cells was performed via hierarchical clustering based on their correlation of global transcriptome and results illustrated four different clusters for ALG (clusters 1 - 4), three clusters for neurons and two additional clusters, one for non-ALG glia (i.e., PNG, SPG, CG, EGN and EGT) and one for a mix of neurons and non-ALG glia (Fig. 1D). These data also illustrated that among all these clusters, astrocytes exhibited the best clustering pattern when they were classified based on their previously-known molecular markers, wherein, their global transcriptome profile was remarkably distinct from neurons and other glia (Fig. 1C and 1D). Differential gene expression analysis was performed between each ALG cluster and the cells of other clusters. Results showed that among all sets of differentially expressed genes (DEGs), the differential expression of 17 genes was common between all ALG clusters, of which eleven genes were enriched and six genes were depleted in ALG (Fig. 1E and 1F). In fact, all the identified ALG clusters (clusters 1 - 4) showed an enriched expression for some previously- known glial markers *CR34335* [46], *Inx2* [47–49] and *Nrv2* [23, 26, 50], and also enrichment for the astrocyte markers *CG9394* [51], *Eaat1* [25, 52, 53], *Gat* [52], *Gs2* [25, 54] and *CG1552* [51] (Fig. 1E), while exhibited a depleted expression for the markers of other cell types (e.g., *gem* and *CG4797* (the PNG markers [25]), *moody* (a SPG marker [25, 46, 55]), *wrapper* (a CG marker [16, 25]) and *CG34340* (a EG marker [16])) (Fig. 1F), supporting the identity of these clusters as astrocytes. The full list of the differentially expressed genes (DEGs) between ALG and non-ALG clusters are presented in supplemental file S1.

### *CG11000*: a gene with enriched expression in Eaat1^+^ cells

As shown above, all the identified ALG clusters were common in expression enrichment of eleven genes, most of which reported as glial or astrocytic markers (Fig. 1E). However, among these genes, the *CR40469* and *Eaat1* genes have been classified by Konstantinides et al. in neuronal clusters of their study (i.e., clusters # 23 and # 43 of *Drosophila* optic lobe) [25], while *Eaat1* predominantly was considered as an astrocyte-specific marker [6, 10, 51, 53, 56, 57] and *CR40469* was not yet reported as a cell type-specific gene in *Drosophila* nervous system.

Sequence analysis of *CR40469* transcript (GenBank accession number: NR_003723.2) demonstrated that it has 96.73% sequence identity with the transcript of the glial marker *CR34335* (GenBank accession number: NR_133508.1) (Fig. 2A), raising the hypothesis that the expression level reported for *CR40469* in the *Drosophila* optic lobe single cells could be influenced by the high-degree sequence homology, hence both transcripts map similarly against *Drosophila* genome leading to highly similar expression counts for both genes. Analysis of the correlation co-efficient between the expression level of these genes across all sequenced single cells of *Drosophila* optic lobe (n = 120,000 cells) showed a remarkable positive correlation (R^2^ = 0.93, *P* < 0.0001, Fig. 2B), strengthening this hypothesis.

**Figure 2.**
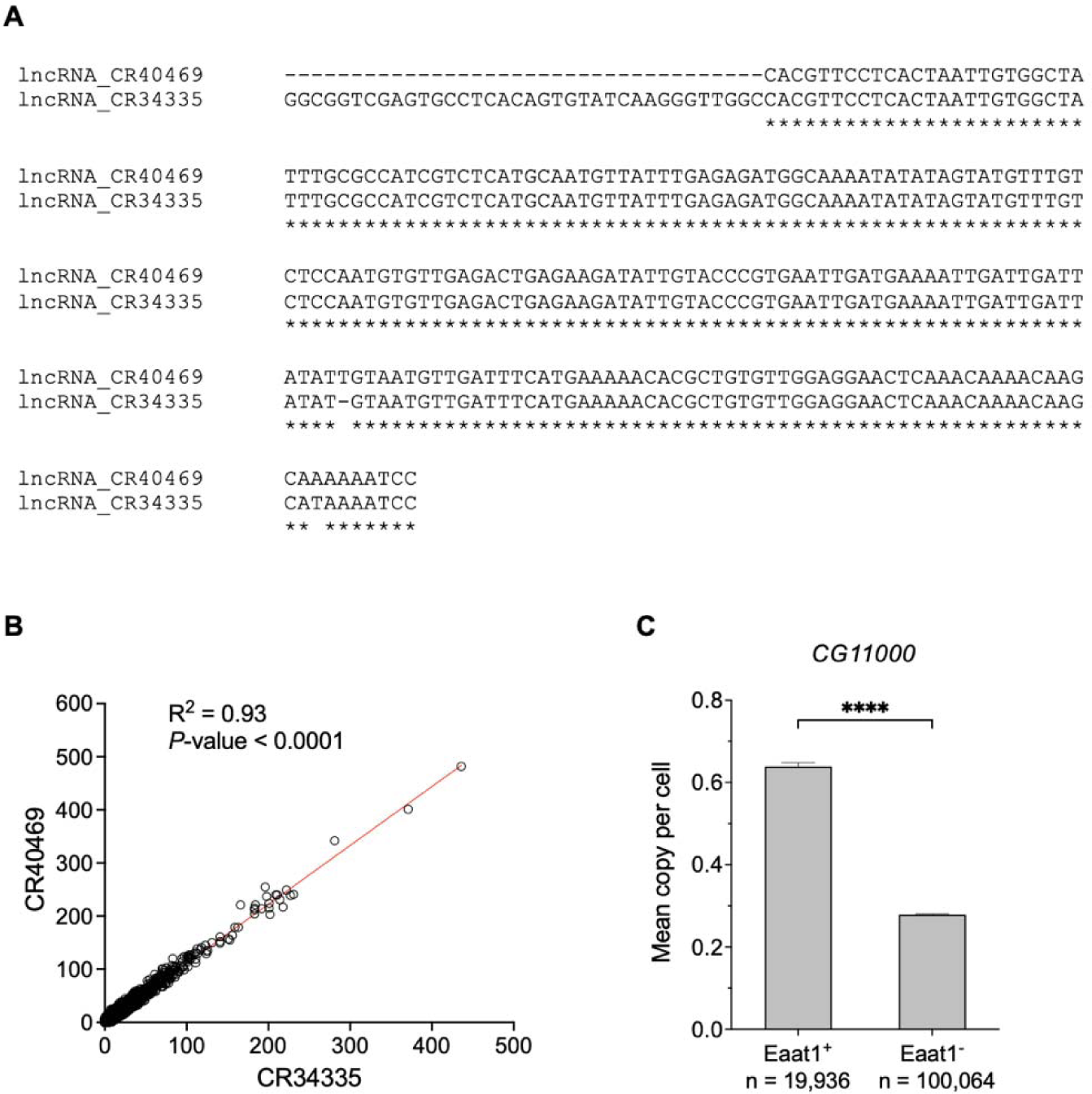
Two candidate genes with differential expression in *Drosophila* astrocytes. **A)** Sequence homology between the transcripts of *CR40469* and *CR34335* genes analyzed by the online tool ClustalW. The nucleotide sequences of both genes are shown in 5’ ➔ 3’ orientation. **B)** Positive expression correlation between *CR40469* and *CR34335* genes in *Drosophila* optic lobe, based on the copy number of each transcript per cells (n = 120,000 single cells). **C)** Increased expression level of *CG11000* gene in *Eaat1*-expressing (Eaat1^+^) cells of *Drosophila* optic lobe. \*\*\*\**P*-value < 0.0001.

*Eaat1* is a known glial marker predominantly expressed in astrocytes [58]. To study which gene(s) are co-expressed with it, first the sequenced single cells of *Drosophila* optic lobe [25] were categorized based on their expression pattern for *Eaat1* gene, then differential gene expression analysis was carried out between the cells ‘with’ and ‘without’ *Eaat1* expression (assigned as Eaat1^+^ and Eaat1^-^ cells, respectively). Results highlighted the differential expression of an uncharacterized gene, named *CG11000*, whose expression was significantly higher in Eaat1^+^ cells (2.43 folds increase, *P*-value < 0.0001) (Fig. 2C), suggesting potential astrocytic function of this gene (similar to *Eaat1* gene). Full list of the upregulated genes in Eaat1^+^ cells (Adjusted *P*-value < 0.01) are presented in the supplemental file S2.

### Positive expression correlation of *CG11000* and *Eaat1* genes in *Drosophila* brain

Given the increased expression level of *CG11000* gene in Eaat1^+^ cells of *Drosophila* optic lobe (Fig. 2C), its expression pattern was also compared to the expression pattern of other known *Drosophila* cell type markers across the full set of single cells in *Drosophila* optic lobe. Results showed a positive expression correlation between *CG11000* and *Eaat1* genes, indicated by co-clustering pattern of these genes based on their similarities in their expression profile across all sequenced single cells in *Drosophila* optic lobe (n = 120,000 cells) (Fig. 3A). Such positive correlation between these genes was also observed across the full set of single cells of *Drosophila* mid-brain (n = 28,695 cells), sequenced by Croset et al. [26] (Fig. 3B).

**Figure 3.**
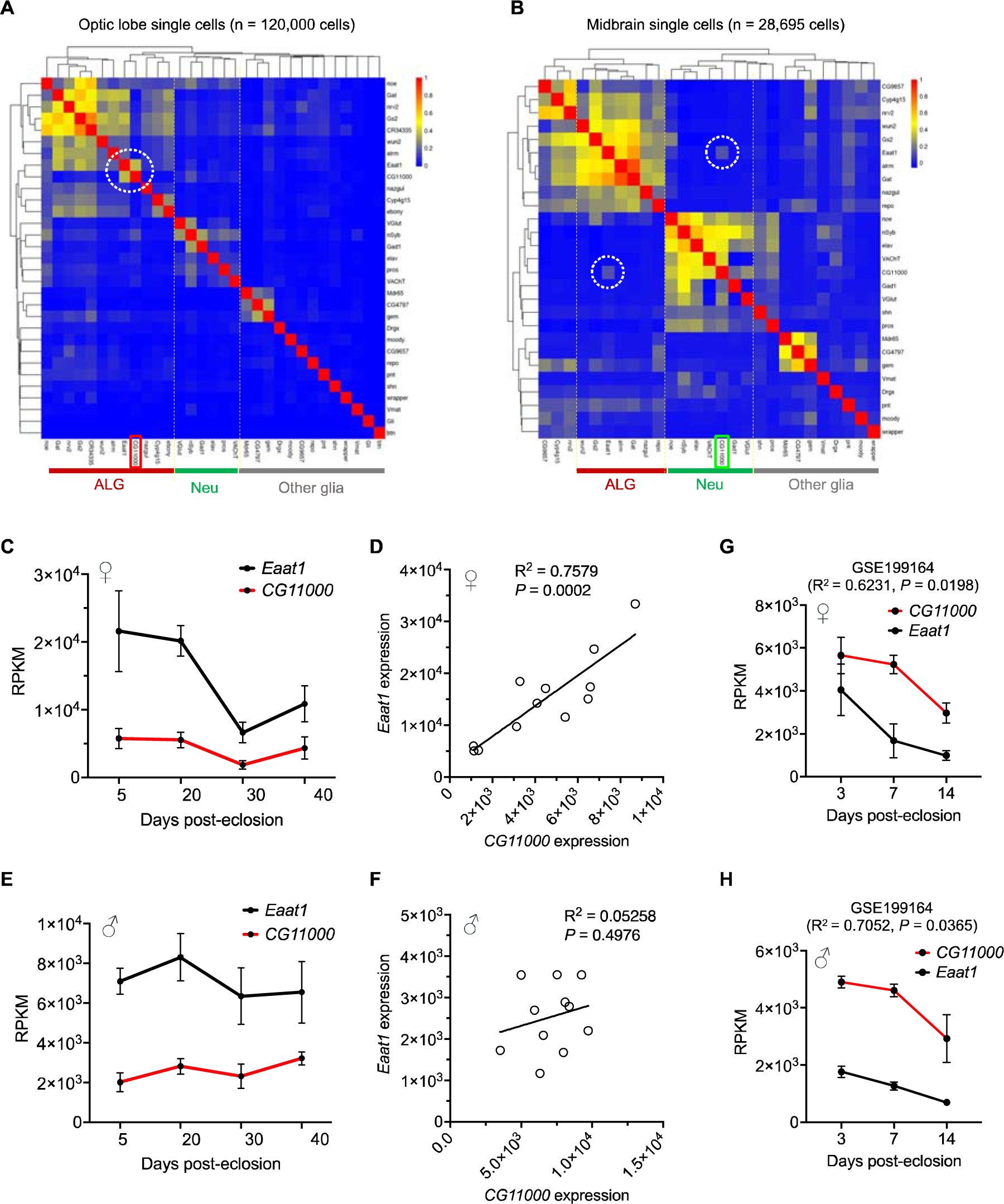
Positive expression correlation of *CG11000* and *Eaat1* genes. **A)** Heatmap analysis of Pearson correlation between *CG11000* gene and neural cell type markers in *Drosophila* optic lobe, using the transcriptome profile of 120,000 sequenced single. *CG11000* is co-clustered with *Eaat1* (dotted circle in the heatmap) due to its similar expression pattern with *Eaat1* expression across all set of single cells in optic lobe. **B)** Heatmap analysis of Pearson correlation between *CG11000* gene and neural cell type markers in *Drosophila* mid- brain, using the transcriptome profile of 28,695 sequenced single cells. Two dotted circles in the heatmap pinpoint the positive expression correlation between *CG11000* and *Eaat1* genes. **C and D)** Correlation analysis of *CG11000* and *Eaat1* expression using a bulk RNA-seq data of developing whole brain from female *D. melanogaster* (n = 12 flies). **E and F)** Correlation analysis of *CG11000* and *Eaat1* expression using a bulk RNA-seq data of developing whole brain from males (n = 11 flies). **G)** Significant expression correlation between *CG11000* and *Eaat1* genes across developmental time-points of female *D. melanogaster*, obtained through bulk RNA-seq data analysis of developing whole brain of the flies. **H)** Significant expression correlation between *CG11000* and *Eaat1* genes across developmental time-points of male *D. melanogaster*, obtained through bulk RNA-seq data analysis of developing whole brain of the flies.

Moreover, the positive expression correlation of *CG11000* and *Eaat1* was further examined in a bulk RNA-seq data of developing *Drosophila* whole brain across different developmental time points of the fly (GSE107049) [27]. Results consistently demonstrated a significant positive correlation between the genes in females (R^2^ = 0.7579, *P*-value = 0.0002) (Fig. 3C and 3D) while at a non-significant level in males (R^2^ = 0.05258, *P*-value = 0.4976) (Fig. 3E and 3F). However, the positive correlation between these genes was evident in both females and males of an additional bulk RNA-seq data from the developing whole brain of *D. melanogaster* (GSE199164) (Fig. 3G and 3H). These data support the positive expression correlation between *CG11000* and *Eaat1*.

### Co-expression of *CG11000* and *Eaat1* genes in *Drosophila* optic lobe

To test whether *CG11000* gene is co-expressed with *Eaat1* in single cells of *Drosophila* optic lobe, the driver lines expressing Gal4 and LexA proteins under the control of *CG11000* and *Eaat1* promoters have been used. For this purpose, the male flies expressing Gal4 under the control of *CG11000* promoter, and expressing LexA under the control of *Eaat1* promoter were crossed to the virgin females transgenic for both UAS-mCD8RFP and LexAop-mCD8GFP constructs (materials and methods, table 1), so that the expression of RFP and GFP in their progenies could reflect the expression pattern of *CG11000* and *Eaat1* genes, respectively. Results illustrated that the RFP signals driven by CG11000-Gal4 were overlapped with the *Eaat1* signals (GFP) in a defined pattern in the *Drosophila* optic lobe in both female (Fig. 4A) and male (Fig. 4B) flies. This data showed the overlapping fluorescence signals of *CG11000* and *Eaat1* genes in a subset of single cells in *Drosophila* optic lobe, which is consistent with the observed positive correlation between the expression of these genes (Fig. 3).

**Figure 4.**
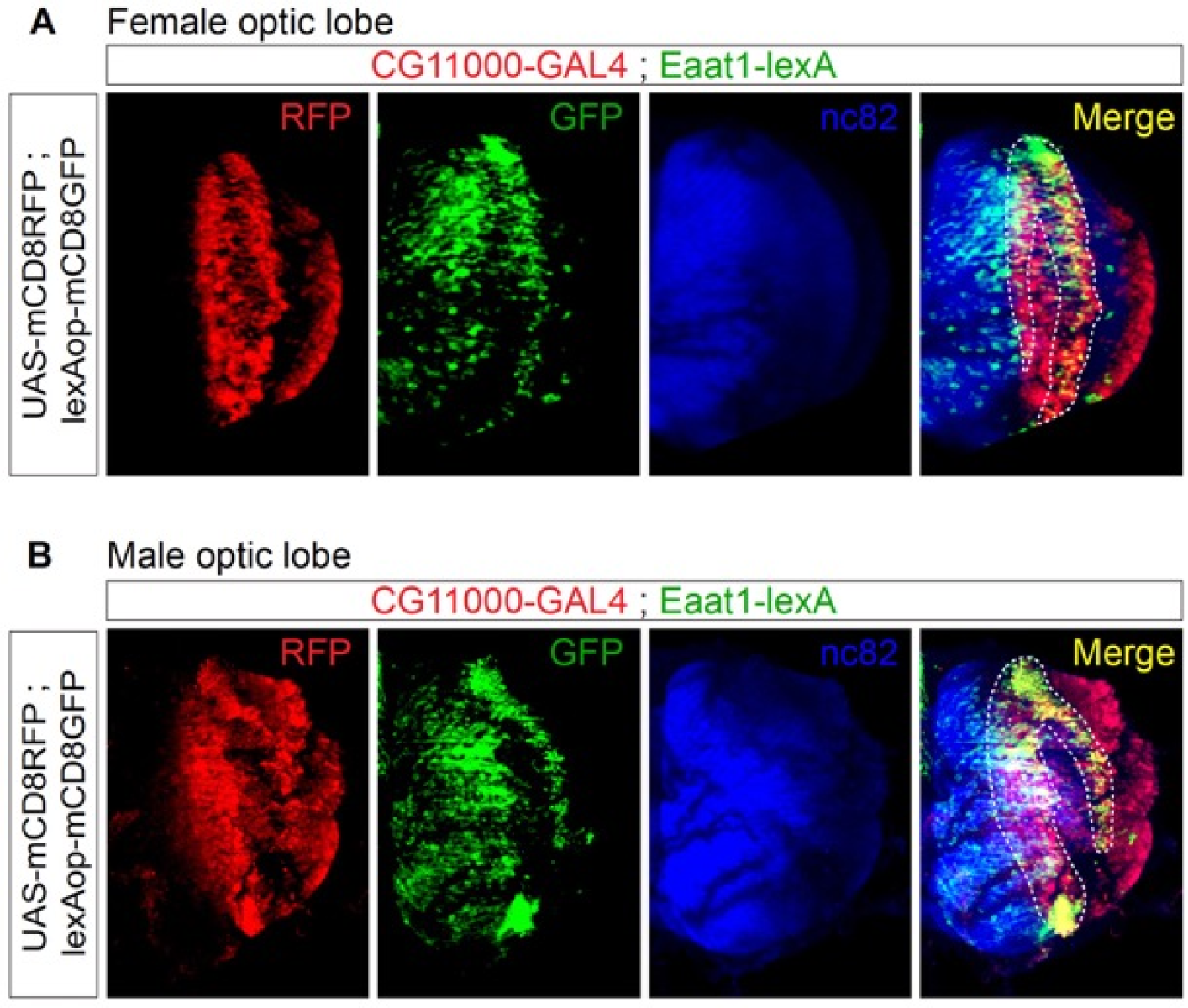
Co-expression analysis of *CG11000* and *Eaat1* in *Drosophila* optic lobe. **A)** Overlapping fluorescence signals correspond to the *CG11000* (RFP; red) and *Eaat1* (GFP; green) expression, tested in the optic lobe of a female *D. melanogaster*. **B)** Overlapping fluorescence signals correspond to the *CG11000* (RFP; red) and *Eaat1* (GFP; green) expression, tested in the optic lobe of a male *D. melanogaster*. The overlapping signals (encircled by dotted lines) represent the cells co-expressing both RFP and GFP which correspond to the expression of *CG11000* and *Eaat1*, respectively. For both immunostaining experiments, an antibody against nc82 was used for labeling the neuropil (blue).

### Sex-biased developmental effects of *CG11000* gene in *D. melanogaster*

For functional analysis of *CG11000* gene, its expression was suppressed using a specific RNAi expressed in both female and male flies in a cell type-specific manner. To this end, virgin females transgenic for the UAS-CG11000 RNAi construct were crossed to a set of male Gal4 driver lines, enabling their progenies to express the *CG11000* RNAi in different cell types of their nervous system (materials and methods; table 2). Also, the flies without any expression of *CG11000* RNAi were used as the control group of the experiment.

Through counting the eclosed flies of the crosses, unexpectedly, we observed a sex-biased eclosion rate for the progenies expressing *CG11000* RNAi, wherein most males expressing *CG11000* RNAi did not develop into adult flies (Fig. 5A), while the females with the same genotypes normally developed into adults (Fig. 5B), suggesting a male-specific developmental function for *CG11000* gene in *D. melanogaster*. In the control group, similar eclosion rate was observed for both male and female progenies.

**Figure 5.**
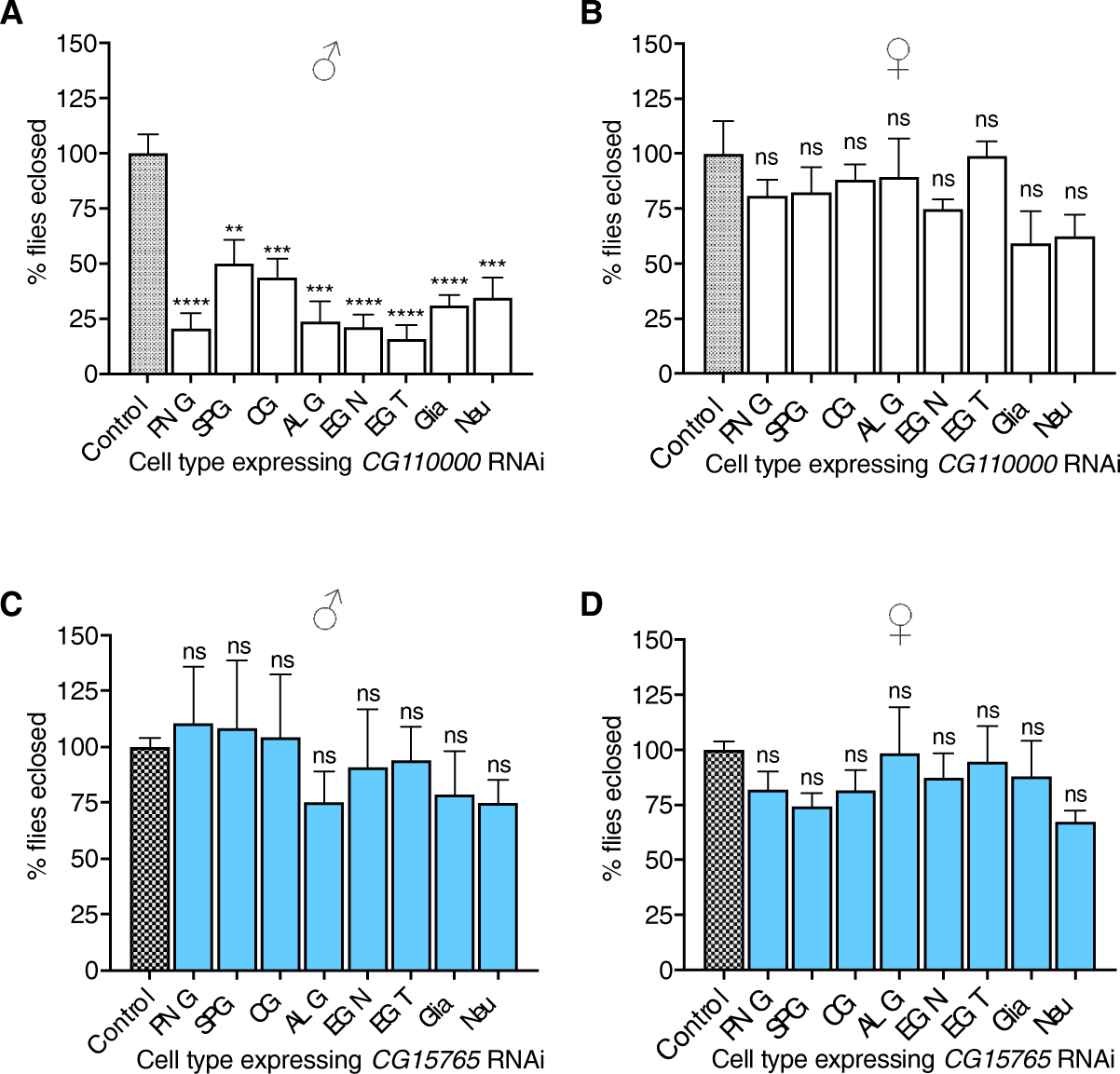
The effects of *CG11000* knock down on normal development of *D. melanogaster*. **A)** In comparison with the control group (lacking expression of *CG11000* RNAi), the majority of male flies with cell type-specific expression of *CG11000* RNAi could not develop into adults, indicated by significant reduced level of eclosion rate (%) in males. **B)** *CG11000* knock down in females, however, did not show any developmental effects, and differences between the eclosion rates (%) of control females (lacking CG11000 RNAi expression) and the flies with cell type-specific expression of *CG11000* RNAi were not statistically significant (denoted by ‘ns’). **C and D)** The results of a parallel experiment using another RNAi (rather than *CG11000* RNAi) as a control. The eclosion rate (%) of all male and female flies in this control experiment did not show any developmental effects, indicated by similar eclosion rate (%) of both male and female flies.

A parallel experiment with the same procedure was carried out, using another *Drosophila* gene named *CG15765*, which we found it as a neuron-specific gene (unpublished data), and the results showed a standard ratio of males over females (∼1) for all progenies expressing *CG15765* RNAi in different neural cell types (Fig. 5C and 5D), supporting the specificity of *CG11000* RNAi to impede development of male progenies.

### *CG11000* affects locomotion and lifespan in *D. melanogaster*

As revealed by this study, the expression levels of *CG11000* and *Eaat1* are positively correlated. On the other hand, *Eaat1* is a well-known astrocytic marker whose expression in astrocytes was shown to be essential for locomotion activities in *D. melanogaster* [53]. To investigate if *CG11000* gene could also affect locomotion in *D. melanogaster*, climbing assay was performed on adult flies expressing the *CG11000* RNAi in their different neural cell types including astrocytes. Results showed that the flies which expressed the *CG11000* RNAi specifically in their ALG had defects in climbing ability (Fig 6A), suggesting that this gene is critical to this cell type. However, ALG-specific knockdown of *CG11000* in female flies did not lead to the above-mentioned defect in climbing activities (Fig 6B), suggesting that the ALG-specific effect of *CG11000* on locomotion behavior in *D. melanogaster* is male-specific. In females, however, slight changes in locomotion activities were identified when *CG11000* gene was supressed in EGN and SPG cells but not ALG cells (Fig 6B). Therefore, astrocytic function of *CG11000* in ALG takes on a great significance only to males (instead of females) in terms of climbing activity.

**Figure 6.**
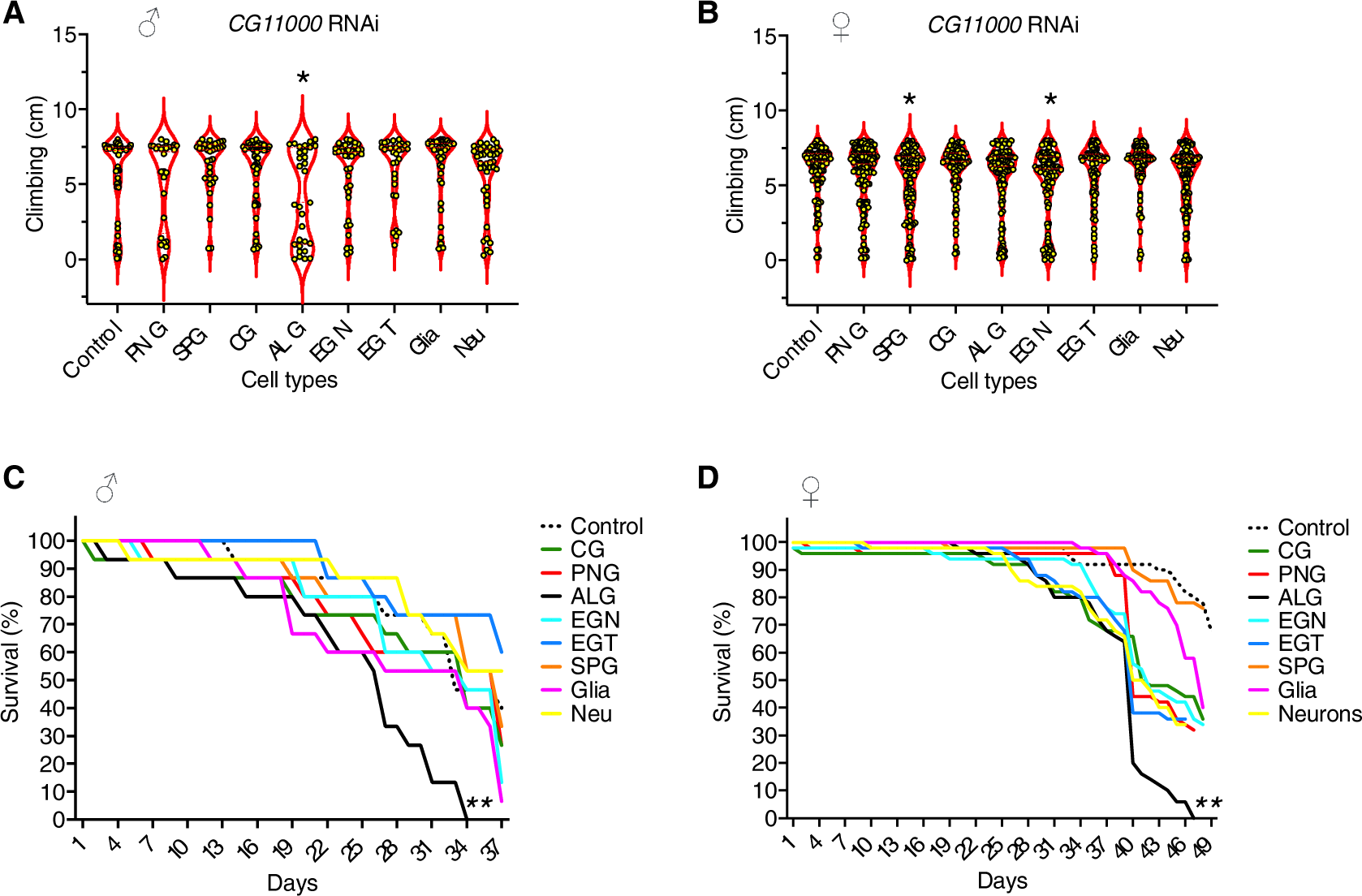
The effects of *CG11000* knock down on locomotion and lifespan of *D. melanogaster*. **A)** ALG-specific effect of *CG11000* expression suppression on climbing activities of male *D. melanogaster* (*P*-value < 0.05). **B)** SPG- and EGN-specific reduced climbing activities in female *D. melanogaster* under expression suppression of *CG11000* (*P*-value < 0.05). **C and D)** ALG-specific reduction of lifespan was observed for both male and female *D. melanogaster*, under expression suppression of *CG11000* gene (*P*-value < 0.01). ALG = astrocyte-like glia, SPG = sub-perineural, EGN = neuropil ensheathing glia.

To test the possible function of *CG11000* gene in *D. melanogaster* lifespan, we supressed its expression in different neural cell types using expressing the *CG11000* RNAi under regulation of cell type-specific Gal4 expression, then their lifespan for 40∼50 days was measured. The most remarkable decreases in the lifespan were identified specifically for the flies expressing the *CG11000* RNAi in their ALG cells (Fig. 6), supporting the astrocytic function of *CG11000* in shortening the lifespan of *D. melanogaster*. The ALG-specific effects of *CG11000* RNAi on lifespan was identified in both male (Fig. 6C) and female flies (Fig. 6D), suggesting that the function of *CG11000* gene on *Drosophila* lifespan is not sexually dimorphic.

## Discussion

Like other metazoans, the *D. melanogaster* brain is formed by a large number of neurons and glia, each with distinct transcriptome profile conferring them unique structural and functional characteristics [46, 59]. Cells do not act individually in brain but instead they make clusters of single cells (known as cell types [10]), each with specialized function and/or organization in different neural circuits of *Drosophila* brain [60–62]. The perineurial glia (PNG), subperineurial glia (SPG), cortex glia (CG), ensheathing glia (EG) and neurons are the main well-characterized cell types in the *Drosophila* nervous system [9, 15, 16, 63, 64]. Moreover, establishing and maintaining the identity of these cell types during development of the fly requires expression of unique combinations of genes, known as their molecular markers [9, 15, 16, 23]. These markers are the genes whose expression implements particular biological processes or signaling pathways to specialize the development and function of a respective cell type [65, 66].

In this study, we used the previously known cell type-specific markers (Fig. 1A) to classify the sequenced single cells of *Drosophila* optic lobe into their corresponding cell types. At least, two different molecular markers were analyzed of each cell type. These markers were among the best well-documented ones in *D. melanogaster*, whose Gal4/UAS driver lines were also frequently used for characterization, mapping and functional analysis of the respective cell types [9, 15, 16, 25, 63, 64]. On the other hand, availability of the transcriptome profile of *Drosophila* brain at single cell level enabled us to reexamine the identity of our classified cells, based on their similarities in their entire transcriptome. As the result, the single cells belonging to the ALG cell type exhibited to have a far distinct transcriptome versus PNG, SPG, CG, EG and neuronal classes when they were classified by expression pattern of five markers (*Eaat1*, *alrm*, *Gs2*, *ebony* and *Gat*) (Fig. 1B and 1C). Differential gene expression analysis showed significant enrichment of 11 genes while depletion of 6 genes in the identified ALG clusters in comparison with other cell types (Fig. 1E and 1F). In addition, while most of these identified ALG-enriched genes were among the previously-reported ALG-enriched/specific markers, the *Eaat1* and *CR40469* genes were determined as neuronal markers in the study by Konstantinides et al. [25] (their supplemental file S1; clusters # 23, 43 and 45). These findings led us to focus on these genes which we found them as the top ALG-enriched genes and also myriad number of previous studies characterized the *Eaat1* as an astrocytic (ALG) marker [10], and no cell type specificity was reported for *CR40469*, yet.

*CR40469* (also known as *lncRNA:CR40469*) spans a region of 213 kb on the X chromosome of *D. melanogaster* (NCBI GenBank, 2023 update). This gene is known to encode a long noncoding RNA with 214 nucleotides but without functional characterization [67–69]. Moreover, due to the high sequence identity rate between *CR40469* gene and the glia marker gene *CG34335*, functional study of *CR40469* via RNAi-based studies will be challenging. However, in a study by Rois et al. [70], a genomic knock out of this gens in *D. melanogaster* was constructed and through differential gene expression analysis the greatest expression level alterations were found for the genes located on the *Drosophila* chromosome X, nearby the *CR40469* locus (e.g., *CG42259*, *png*, and *CG4313* genes). According to this data, a trans-acting function was attributed to *CR40469* long noncoding RNA, by which it regulates the expression of its neighboring genes on chromosome X [70]. Here we found that the expression pattern of *CR40469* across the single cells of *Drosophila* optic lobe resembled that of *CR34335* gene which was previously reported as a glial marker gene [46] which could be in part due to the high-degree homology rate between these transcripts (Fig. 2A and 2B). However, further functional studies are required to elucidate the role of *CR40469* in *Drosophila* nervous system. No homologue of this gene was found in the human genome.

Consistent with our results for ALG-enrichment of *CG40469* gene, it was found by Croset et al. [26] as glial/astrocytic marker, suggesting the pivotal role of *CG40469* gene in these cell types.

Unlike *CR40469* gene, *Eaat1* is a well-characterized gene crucial to development, physiology and behaviors of *D. melanogaster*. When we categorized the sequence single cells of *Drosophila* optic lobe based on their expression pattern for *Eaat1*, we found the differential expression of an uncharacterized gene, named *CG11000*, which was significantly higher in Eaat1^+^ cells versus the single cells without *Eaat1* expression (i.e., Eaat1^-^ cells). This preliminary data compelled us to investigate the expression correlation of *Eaat1* and *CG11000* genes in *Drosophila* brain. Interestingly, the positive correlation was existed between these genes in total set of single cells in *Drosophila* optic lobe and mid-brain, as well as their positive expression correlation in developing *Drosophila* whole brain (Fig. 3). Beside positive correlation, *CG11000* was clustered in the ALG cells of Drosophila optic lobe (Fig. 3A), while clustered in the neuronal cluster in the mid-brain (Fig. 3B) suggesting some non-ALG expression pattern for this gene. Moreover, the observed positive correlation between *CG11000* and *Eaat1* genes might be due to the high frequency of the cells co-expressing them. In support of this expression patterns, the expression pattern of these genes were investigated in single cell collection of adult *Drosophila* whole brain from SCope database [71] (R^2^ = 0.6611, *P*-value <0.0001; Supplemental file S3).

Our data also provides a comparative assessment between two approaches for cell type identification: 1) cell type identification based on the pre-defined set of molecular markers, and 2) cell type identification based on the similarities in whole transcriptome profile.

Since a set of multiple marker genes could not represent the complete genetic profile of a respective cell type, and on the other hand, different cell types may have similar transcriptome profile but different only in the expression of a limited number of genes, each of the above-mentioned approaches has its own pros and cons that should be considered for cell type identification purposes. For instance, *Eaat1* expression pattern *per se* could not distinguish ALG cells among other single cells because *Eaat1* is also expressed at low levels in a subset of neurons [25]. Similarly, some clusters in the Konstantinides et al. study were identified as neurons based on their entire transcriptome profile (clusters # 4 and 14) while they were positive for the glial marker *repo* [25] which necessitate using additional markers on these clusters to reveal their actual identity. Also, studies have shown that while Repo protein is a good marker for glial cells [72, 73], its transcripts are lowly expressed and often not detected in glial clusters [74], which challenges the use of *repo* gene as a glial marker in RNA-seq data analyses.

In Drosophila research, the binary systems such as Gal4/UAS and LexA/lexAop are powerful genetic tools to map the expression pattern of the genes to different neural cell types [14]. here, we used these systems to trace the distribution of mCD8RFP and mCD8GFP signals across the *Drosophila* optic lobe driven by the *CG11000* and *Eaat1* promoters, respectively. The overlapped fluorescent signals of these genes illustrated their co-expression in a set of single cells in

Drosophila optic lobe (Fig. 4). However, some single cells in *Drosophila* optic lobe did not show such co-expression pattern, particularly the cells in the cortical region of optic lobe which were positive for *CG11000* while negative for *Eaat1* (Fig. 4). This data is also consistent with the observed expression pattern of the *CG11000* and *Eaat1* genes in a cluster of single cells in adult Drosophila brain obtained from the SCope, wherein some single cells express *CG11000* transcripts while negative for *Eaat1* expression (Supplemental file S3).

In addition to the expression pattern analysis of *CG11000* gene, its functional analysis was performed through cell type-specific expression of an RNAi against this gene in male and female flies.

Unexpectedly, through collecting and counting the progenies with *CG11000* suppression by RNAi, we observed a reduced eclosion rate exclusively in male progenies (but not in females), suggesting a male-specific developmental function for *CG11000* gene in *D. melanogaster*.

However, RNAi-mediated knock down of *CG11000* in adult flies caused ALG-specific defects in climbing activities of male flies (but not females), while such defects in the climbing activities of female flies were found to be SPG- and EGN-specific. Such sex-biased phenotype changes under CG11000 knock down could suggest that CG11000 exerts its biological functions through cellular and/or molecular pathways which differ in males and females, while the expression pattern of *CG11000* is similar in both sexes (Fig. 4). However, further functional studies are required to elucidate the molecular mechanisms underlying the sex-biased effects of *CG11000* in the development and physiology of *D. melanogaster*.

The analysis of lifespan in the flies expressing *CG11000* RNAi in different cell types revealed the greatest effect when *CG11000* was suppressed specifically in ALG cells of both sexes, thus verifying the functional specificity of this gene to ALG cell type.

Moreover, CG11000 knock down in different cell types of adult flies, resulted in shortened lifespan in both males and females in an ALG-specific manner. This result highlights the crucial role of CG11000 gene in normal lifespan of D. melanogaster which is not sexually dimorphic.

## Supporting information

Supplemental file previous S1

## Acknowledgments

The stocks used in this study were from the Bloomington Drosophila Stock Center (NIH P40OD018537). We thank Khoi-Nguyen Ha Nguyen for his assistance in the experiments performed in this study.

