## Supplemental file S3 for "The astrocyte-enriched gene *CG11000* plays a crucial role in the development, locomotion and lifespan of *D. melanogaster*"

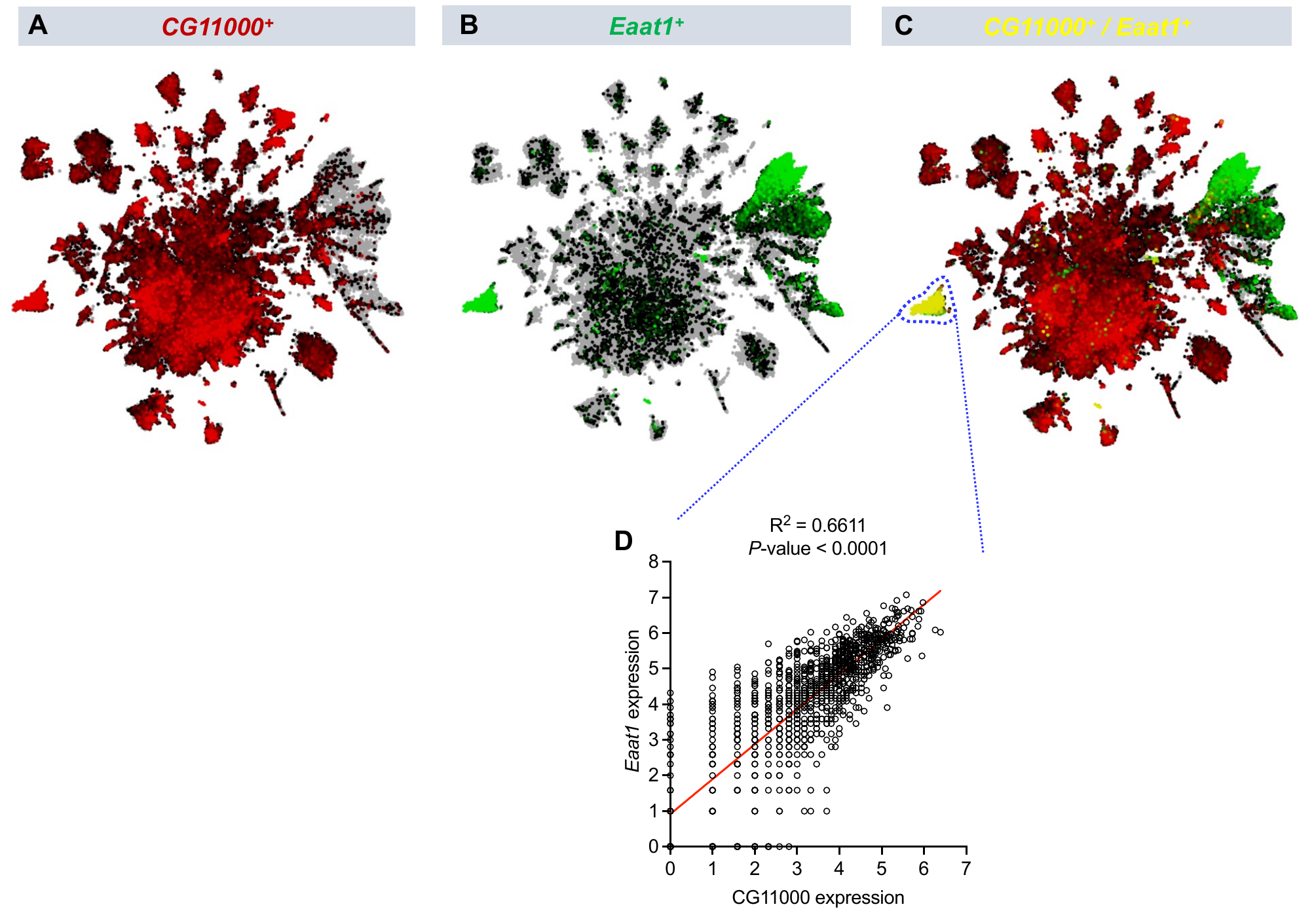

**Supplemental file S3. Co-expression of *CG11000* and *Eaat1* genes in a set of single cells in adult *Drosophila* brain adapted from the Scope database.**

**A)** Expression pattern of *CG11000* transcripts.

**B)** Expression pattern of *Eaat1* transcripts.

**C)** Merged pattern of *CG11000* and *Eaat1* expression across the single cells co-expressing them. **D)** Correlation analysis of the expression level of *CG11000* and *Eaat1* genes.
